# Biophysical Constraints Dictate the Stability of Social Traits in *Pseudomonas aeruginosa* Aggregates

**DOI:** 10.64898/2026.03.18.712675

**Authors:** Kathleen O’Connor, Nicholas Larson, Sheyda Azimi, Stephen P. Diggle

## Abstract

The maintenance of cooperative behaviors within microbial populations remains an evolutionary puzzle, particularly in spatially structured environments like those found in chronic infection. In *Pseudomonas aeruginosa*, quorum sensing (QS) directs the production of costly public goods that are vulnerable to exploitation by non-producing cheaters. However, the biophysical mechanisms that stabilize this cooperation in complex environments remain poorly understood. Here, we elucidate how the biophysical properties of the bacterial cell surface govern the micron-scale architecture of bacterial aggregates to promote or constrain the stability of cooperation. Using a polymer-structured growth environment, we demonstrate that cell surface hydrophobicity acts as a primary determinant of spatial organization, where hydrophilic wild-type cells assemble into “stacked” aggregates, whereas hydrophobic O-specific antigen-deficient cells form dense “clumps”. In mixed populations, distinct surface properties exhibit phase-separation-like immiscibility, segregating the cells at the micron scale. We show that this segregation directly suppresses cheater fitness by physically sequestering cooperative clusters. Specifically, hydrophobic QS-deficient cells were effectively excluded by hydrophilic QS cooperators, limiting their access to public goods. Furthermore, invasion experiments revealed that hydrophilic cells are inherently fitter, capable of unidirectionally invading populations with a hydrophobic cell surface and independent of social dynamics. Conversely, the need to gain access to nutrient resources can fine-tune these barriers, enabling hydrophilic QS-deficient cells to cluster proximally to hydrophobic cooperators. These findings establish that bacterial cell surface traits impact strongly on microbial social interactions and offer new insights into the maintenance and loss of cooperative traits in chronic infections such as those found in chronic lung infections.

**Significance Statement:** Cooperative behaviors are essential for the survival of pathogenic bacteria like *Pseudomonas aeruginosa*, yet they are continually threatened by “cheater” cells that exploit shared public good resources. We demonstrate that the physical properties of the cell surface, specifically hydrophobicity, act as a key architect of microbial social structure. In polymer-rich environments, differences in surface hydrophobicity drive a distinct “oil-and-water” segregation between cell populations. This micron-scale phase separation physically sequesters cooperative cells, insulating them from exploitation by cheaters. Consequently, spatial structure serves as a biophysical stabilizer of cooperation, limiting the metabolic advantages typically held by cheaters. Our findings demonstrate that microbial social evolution is governed not only by genetic strategies, but by fundamental biophysical constraints that dictate the spatial limits of exploitation.

## Introduction

The evolution of social traits and cooperative behaviors has long fascinated researchers, as cooperation is often important for survival under both biotic and abiotic stress (1-4). In bacterial populations, cooperative cells produce “public or common goods”, such as extracellular enzymes, that enhance group survival under stressful conditions like nutrient limitation or predation (5-8). Cheater cells avoid the metabolic costs of producing these goods and instead redirect resources toward growth, gaining a fitness advantage in mixed populations. This raises a central question: how is cooperation maintained despite the continual emergence of cheaters? In animals, cooperation is stabilized by mechanisms such as kin discrimination, where individuals preferentially direct cooperation towards other cooperators (9, 10). Similarly, bacteria employ strategies to restrict cheater access to shared resources. Spatial structuring is one key mechanism that can maintain cooperation, often through kin selection, by physically separating cheaters from cooperators and limiting exploitation (11-15). However, specific spatial structures and scales needed to protect cooperative bacteria remain poorly defined.

The opportunistic pathogen *Pseudomonas aeruginosa* provides a powerful model organism to study these dynamics (16, 17). This bacterium exhibits numerous social behaviors, including quorum sensing (QS), a chemical communication system that regulates the production of extracellular enzymes and other public goods (18-20). While most prior studies have examined QS-mediated interactions in well-mixed cultures, where cheaters and cooperators have equal access to shared public goods (5, 7, 21-30), most bacterial communities reside in spatially structured biofilms or free-floating aggregates (31-35). In such environments, cooperation is predicted to be more stable, likely because spatial proximity limits the spread of cheating cells (11, 36-38). Despite this, the influence of micron-scale organization on cooperation within suspended or surface-associated aggregates, such as those found in cystic fibrosis (CF) chronic lung infections, has not previously been elucidated.

Several biophysical factors, including extracellular DNA (eDNA), exopolysaccharides (EPS), mucin, and cell surface hydrophobicity, influence *P. aeruginosa* aggregate formation (39-43). We previously demonstrated that the loss of O-specific antigen (OSA) glycan moieties in lipopolysaccharide (LPS) alters aggregate assembly in a synthetic CF sputum medium (SCFM2) (42, 44). Mutations in genes involved in OSA biosynthesis and QS regulation are common in chronic infections (45-52), but their combined effects on social dynamics have not been studied to date. Here, we examine how variation in cell surface hydrophobicity and the spatial organization of cells shapes the social interactions in *P. aeruginosa*. Using a spatially structured model that promotes aggregate formation, we manipulated hydrophobicity to control aggregate structure and cell proximity and studied its impact on QS-dependent cooperation and cheating. QS-deficient mutants (Δ*lasR*Δ*rhlR*), which cannot produce exoproteases, rely on public goods generated by wild-type cooperators and thus serve as model cheaters in environments where QS is required for growth (7, 18, 27).

Through three-dimensional confocal microscopy and quantitative image analysis, we show that cells with hydrophilic and hydrophobic surfaces segregate into distinct aggregates, like the immiscibility of oil and water. This segregation at micron-length scales, highlights how spatial organization influences bacterial social interactions. Overall, our findings demonstrate that (i) cheater fitness depends on spatial proximity to cooperators, and (ii) in mixed populations, cell surface hydrophobicity directly influences both the spatial organization of cells and their relative fitness. These results advance our understanding of how cooperation is maintained in spatially-organized bacterial populations and have implications for chronic infections, where *P. aeruginosa* forms persistent aggregates which are often tolerant to antibiotics.

## Results

### Cell Surface Hydrophobicity Dictates Aggregate Architecture and Micron-Scale Variant Immiscibility

Previously we showed that hydrophobicity determines *P. aeruginosa* aggregate-type in SCFM2, which recapitulates the physiological conditions found in CF sputum and includes the addition of mucin and eDNA (42). In order to investigate the robustness of this hydrophobicity-mediated aggregation, we sought to replicate these phenotypes in a more defined environment before assessing the mixing dynamics of different surface types. We therefore assessed aggregation in monocultures using QSM2 (a structured environment where cells require QS activity for growth) containing a proteinaceous carbon source and 0.6 mg/mL eDNA. As expected, hydrophilic wild-type PAO1 formed “stacked” aggregates (Fig. 1A), whereas deletion of the *ssg* gene (PAO1Δ*ssg*), which disrupts O-specific antigen (OSA) assembly and increases cell surface hydrophobicity, lead to the formation of “clumped” aggregates (Fig. 1B), consistent with our prior observations in SCFM2 (42). The aggregation types were independent of fluorescent labels (Fig. S1A,B). To test whether these aggregation rules generalize to conditions where growth is decoupled from social traits, we switched to a non-social growth environment to focus strictly on population mixing behavior across surface types. We combined hydrophilic PAO1 and hydrophobic PAO1Δ*ssg* cells at a 1:1 ratio in a structured environment where growth does not require QS (OS-CAA + 0.6 mg/mL eDNA). After 22–24 h of static incubation, cells with similar surface properties co-aggregated (Fig. 1C, D), whereas in hydrophilic– hydrophobic mixtures, cells segregated into distinct stacked and clumped aggregates (Fig. 1E).

**Figure 1.**
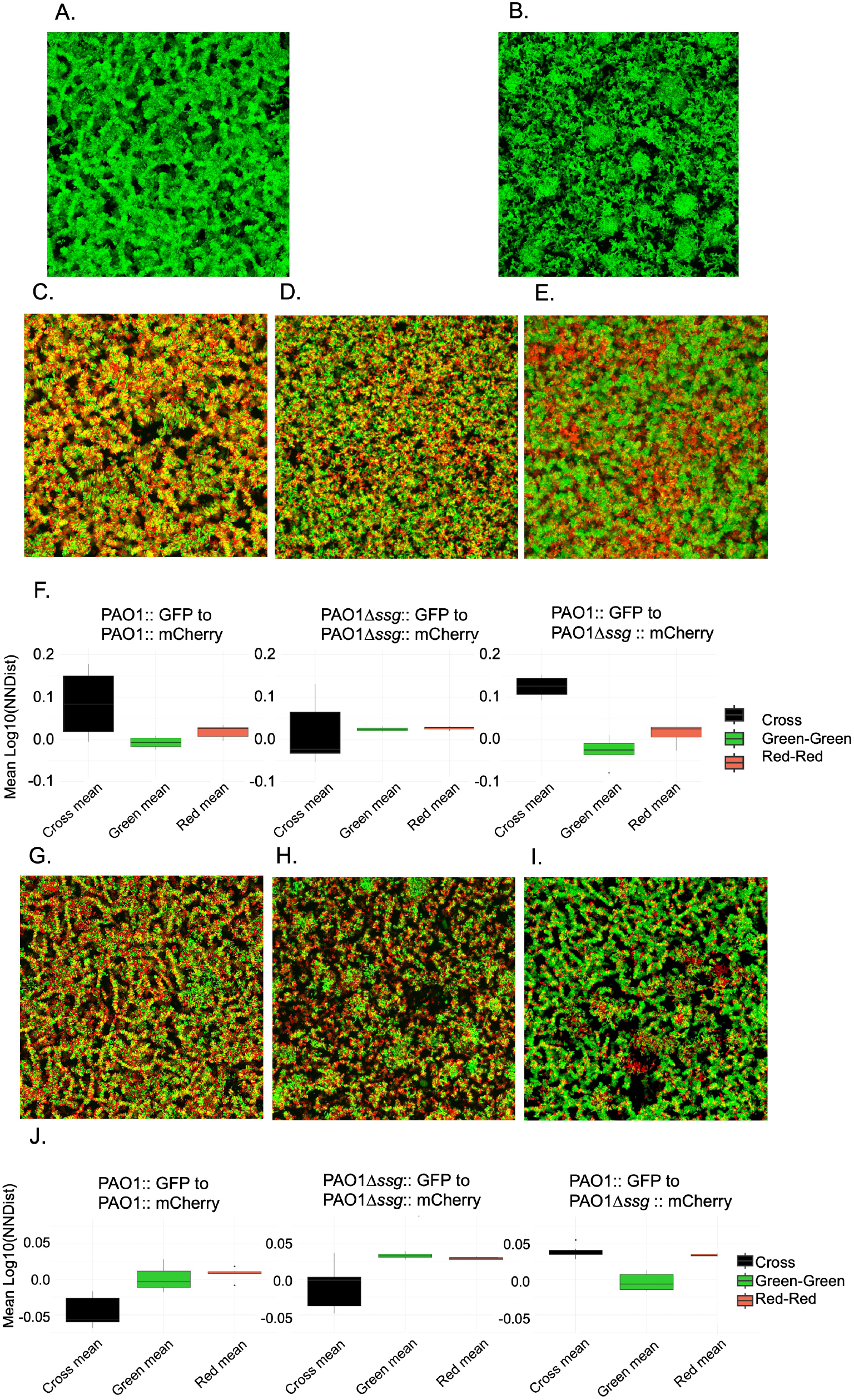
*P. aeruginosa* aggregate assembly and spatial organization is shaped by relative cell surface hydrophobicity. Monocultures of PAO1 (A) and PAO1Δ*ssg* (B) form stacked and clumped aggregates in QSM2. 50/50 mix experiments to determine spatial mixing in asocial media (OS-CAA) using PAO1 cells (GFP and mCherry) (C), PAO1Δ*ssg* cells (GFP and mCherry) (D), and a PAO1::GFP + PAO1Δ*ssg*::mCherry mix (E). (F) Bar plots of Log10 transformed mean nearest neighbor distance (NNDist). (G–I) Similar to (C–E) but performed in social media (QSM2). (J) Bar plots of mean NNDists. Images are 10 µm thick confocal Z-stacks.

We next quantified spatial organization in asocial OS-CAA using a nearest-neighbor distance (NNDist) analysis. We found that cells with different surface properties were significantly farther apart than cells sharing the same surface type, when distances between PAO1 to PAO1 (red-to-red), PAO1Δ*ssg* to PAO1Δ*ssg* (green-to-green), and PAO1 to PAO1Δ*ssg* (cross, red-to-green) were measured (Fig. 1F). In populations of mixed hydrophilic–hydrophobic cells (PAO1::GFP + PAO1Δ*ssg*: mCherry), the percentage of the cross-NNDist area under the curve (AUC) exceeding the same-color distributions was 34.06%, compared to 22.78% for PAO1 and 17.77% for PAO1Δ*ssg* monocultures, indicating greater cell segregation for the mixed hydrophilic-hydrophobic populations (Fig. S1C-E). In homogeneous populations, cross-channel distances were not different from same-color distances or differed only minimally. In contrast, in mixed populations of hydrophilic–hydrophobic cells, cross-NNDist values were significantly elevated with a large fold change (Adj.p < 0.05, Fig. 1F; Table S1-A). Together, these results demonstrate that cell-surface hydrophobicity is a dominant determinant of aggregate architecture and strain mixing, driving micron-scale segregation that likely shapes social interactions and access to public goods in structured environments.

We next determined whether the social requirements of the environment impact how cells cluster. Similar to our findings in the asocial OS-CAA media, in our structured social media (QSM2), cells of the same cell surface co-aggregated while cells with different cell surfaces had more discrete clustering by cell surface type, demonstrating that aggregate-type and variant mixing is similar in social and asocial medias (Fig. 1G-I). Similarly to what was observed in OS-CAA, in QSM2, the highest NNDist AUCs was for mixed populations of hydrophobic-hydrophilic cells (16.71%) (Fig. S2A-C). For mixed populations of strains with the same cell surface properties, there were generally no differences in spatial organization and proximity (Fig. 1J; Table S1-B), and only distances between PAO1 and PAO1Δ*ssg* cells in mixed populations was there a significant difference (Cross v Green, Adj.p < 0.05, Table S1-B). Unlike what was observed in asocial media, in QSM2, the hydrophilic PAO1 cells clustered together, while hydrophobic cells (PAO1Δ*ssg*) were as likely to cluster with either hydrophilic or hydrophobic cells.

### Surface Hydrophobicity Determines Aggregation Patterns Between QS^+^ And QS^−^ Cells in Asocial Environments

After determining that the different environments resulted in changes of cell clustering in mixed populations of hydrophilic and hydrophobic cells, we next studied the impact on a population of mixed QS-cooperating PAO1 cells (QS^+^) and non-cooperating PAO1Δ*lasR*Δ*rhlR* (QS^−^) cells in OS-CAA (asocial environment). We found that QS does not play a role in aggregate-type; specifically, PAO1Δ*lasR*Δ*rhlR* or PAO1Δ*ssg*Δ*lasR*Δ*rhlR* formed stacked or clumped aggregates respectively in OS-CAA, and when QS^+^ and QS^−^ cells of the same surface were mixed, they co-aggregated and formed mixed stacked or clumped aggregates (Fig. 2A,B), while mixed populations of QS^+^ and QS^−^ cells with different cell surface hydrophobicity levels resulted in discrete aggregates composed mostly of either red or green cells (Fig. 2C,D). In mixed populations of cells with the same cell surface properties, there was nearly no cross NNDist AUC above green and red NNDist AUCs (Fig. S3A,B), indicating that QS^+^ green cells were as likely to cluster with red QS^−^ cells. When hydrophilic QS^+^ cells were mixed with hydrophobic QS^−^ cells, there was a large cross NNDist AUC that was above the red and green NNDist AUCs (27.45%, Fig. S3C). Surprisingly, for mixed populations of hydrophobic QS^+^ and hydrophilic QS^-^ cells, there was no cross AUC greater than red or green AUCs (Fig. S3D). Additionally, in the hydrophilic QS^+^; hydrophobic QS^-^ mixed population, there was a noticeable green peak at a low NNDist value to the right of main peak, suggesting that hydrophilic QS^+^ cells cluster especially close to the same cell types in the presence of hydrophobic QS^−^ cells (Fig. S3C).

**Figure 2.**
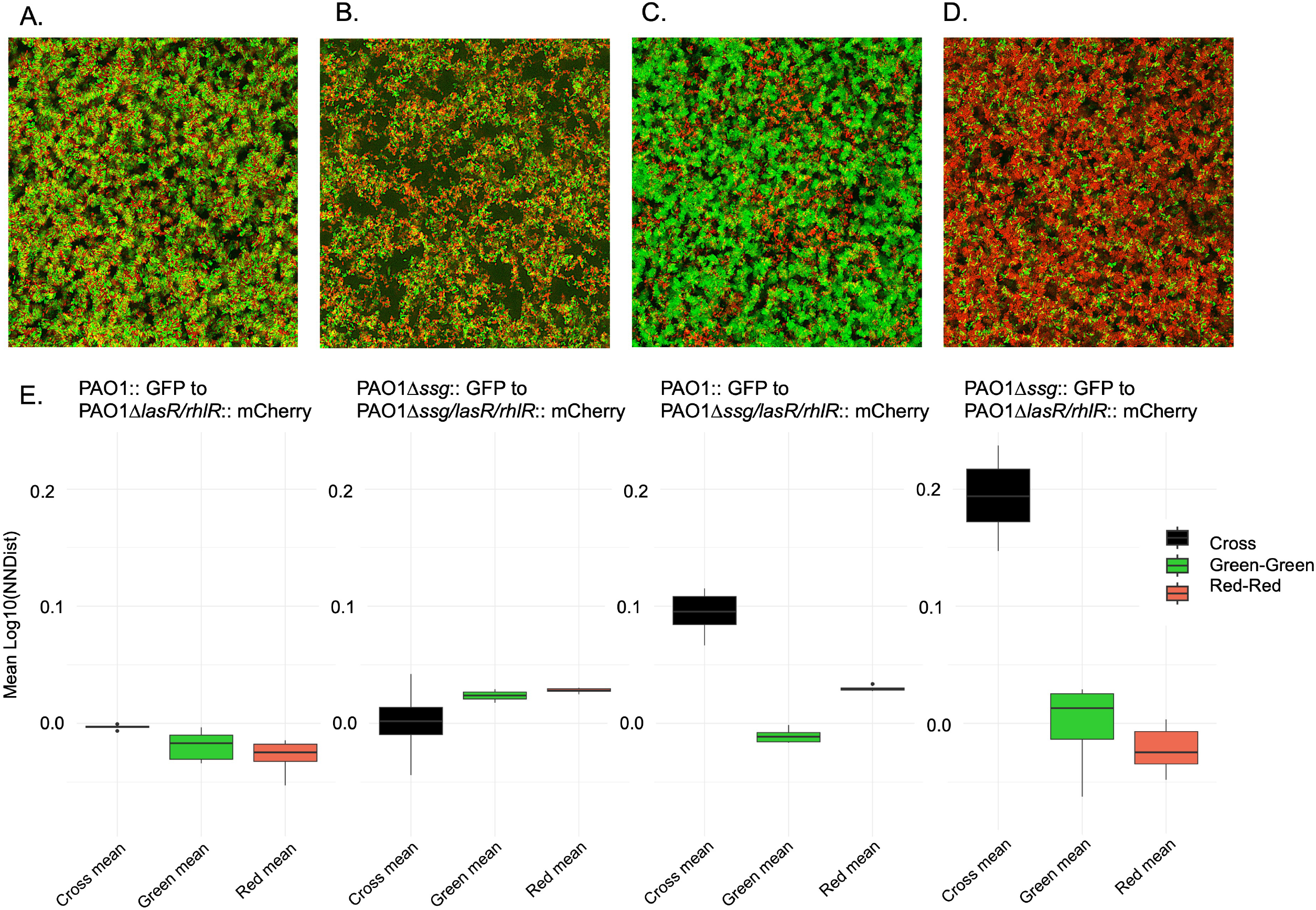
Cell surface hydrophobicity determines the spatial organization of mixed *P. aeruginosa* populations in asocial environments. QS^+^ and QS^-^ strains with varying hydrophobicity were grown in asocial medium (OS-CAA) to assess mixing independent of social cheating. (A–D) Confocal images of 50:50 mixed populations inoculated at OD 0.01: (A) PAO1::GFP+ PAO1Δ*lasR*Δ*rhlR*::mCherry; (B) PAO1Δ*ssg*::GFP + PAO1Δ*ssg*Δ*lasR*Δ*rhlR*::mCherry; (C) PAO1::GFP + PAO1Δ*ssg*Δ*lasR*Δ*rhlR*::mCherry; and (D) PAO1Δ*ssg*::GFP + PAO1Δ*lasR*Δ*rhlR*::mCherry. (E) Log10 transformed NNDist per image. Higher cross-NNDists (black) relative to self-NNDists (green/red) in mixed-hydrophobicity populations indicate decreased mixing compared to populations with matched hydrophobicity.

For populations with mixed hydrophilic QS^+^ and QS^−^ cells (PAO1::GFP and PAO1Δ*lasR*Δ*rhlR*::mCherry), the cross NNDists were significantly higher than the mean NNDists for green and red, although, all values were similar, with low fold changes (1.48 and 1.3 accordingly) (Fig. 2E, Table S1-C). For populations with mixed hydrophobic QS^+^ and QS^−^ cells (PAO1Δ*ssg*::GFP; PAO1Δ*ssg*Δ*lasR*Δ*rhlR*::mCherry), NNDists between all cells were similar and cross NNDists between QS^+^ and QS^−^ cells was not significantly different from red or green NNDists (Fig. 2E, Table S1-C). For mixed populations of hydrophobic and hydrophilic cells (PAO1::GFP; PAO1Δ*ssg*Δ*lasR*Δ*rhlR*::mCherry and PAO1Δ*ssg*::GFP; PAO1Δ*lasR*Δ*rhlR*::mCherry), the cross NNDist was larger compared to red and green NNDists (Fig. 2E, Table S1-C), suggesting that QS^+^ and QS^-^ cells with similar surface properties cluster closely to each other. Taken together, these data demonstrate that in an asocial environment, surface-driven physical segregation, rather than QS social-profile, dictates how closely strains interact.

### Social Requirements in QSM2 Restricts the Growth and Spatial Integration Of QS^−^ Cells

We next tested the spatial organization of mixed populations in QSM2, where QS activation is required for survival. We first demonstrated that QS^+^ cells (PAO1; PAO1Δ*ssg*) had a higher fitness than QS^−^ cells (PAO1Δ*lasR*Δ*rhlR*; PAO1Δ*ssg*Δ*lasR*Δ*rhlR*) when grown in monoculture in QSM2 (Fig. 3A-D), where QS^−^ strains could be rescued with the addition of exogenous protease in the form of proteinase K (PK) (Fig. S4A) due to the breakdown of the proteinaceous carbon source in this media. In mixed populations of cells with the same surface properties, QS^+^ and QS^−^ cells co-aggregated (Fig. 3E,F). However, we found that, hydrophobic QS^−^ cells do not aggregate with hydrophilic QS^+^ cells and instead form distinct aggregates (Fig. 3G). In contrast, hydrophilic QS^−^ cells do co-aggregate with hydrophobic QS^+^ cells, which is in line with findings in asocial media (Fig. 4H).

**Figure 3.**
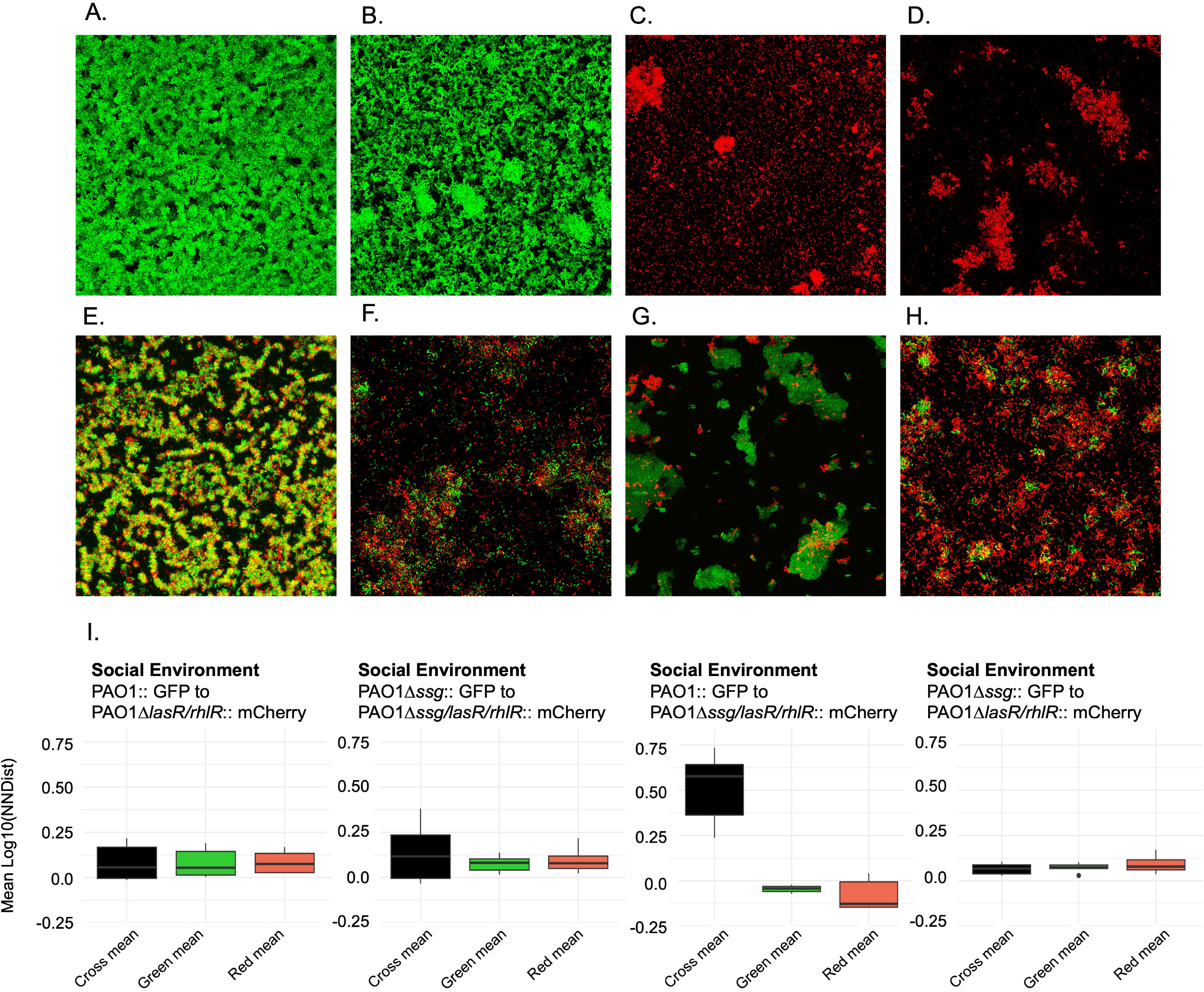
Social environmental requirements modulate aggregate spatial organization and interactions between QS^+^ and QS^-^ strains. (A–D) Growth of QS^+^ PAO1 (A) and PAO1Δ*ssg* (B) strains and QS^-^ PAO1Δ*lasR*Δ*rhlR* (C) and PAO1Δ*ssg*Δ*lasR*Δ*rhlR* (D) strains in QSM2. QS^+^ strains exhibit significantly higher growth than QS^-^ strains (Holm-adjusted t-test; adj. p < 0.001). (E–H) Confocal images of 50:50 QS^+^: QS^-^ mixes in QSM2: (E) PAO1::GFP + PAO1Δ*lasR*Δ*rhlR*::mCherry; (F) PAO1Δ*ssg*::GFP + PAO1Δ*ssg*Δ*lasR*Δ*rhlR*::mCherry; (G) PAO1::GFP + PAO1Δ*ssg*Δ*lasR*Δ*rhlR*::mCherry; and (H) PAO1Δ*ssg*::GFP + PAO1Δ*lasR*v*rhlR*::mCherry. (I) Mean Log10 transformed NNDist per image.

**Figure 4.**
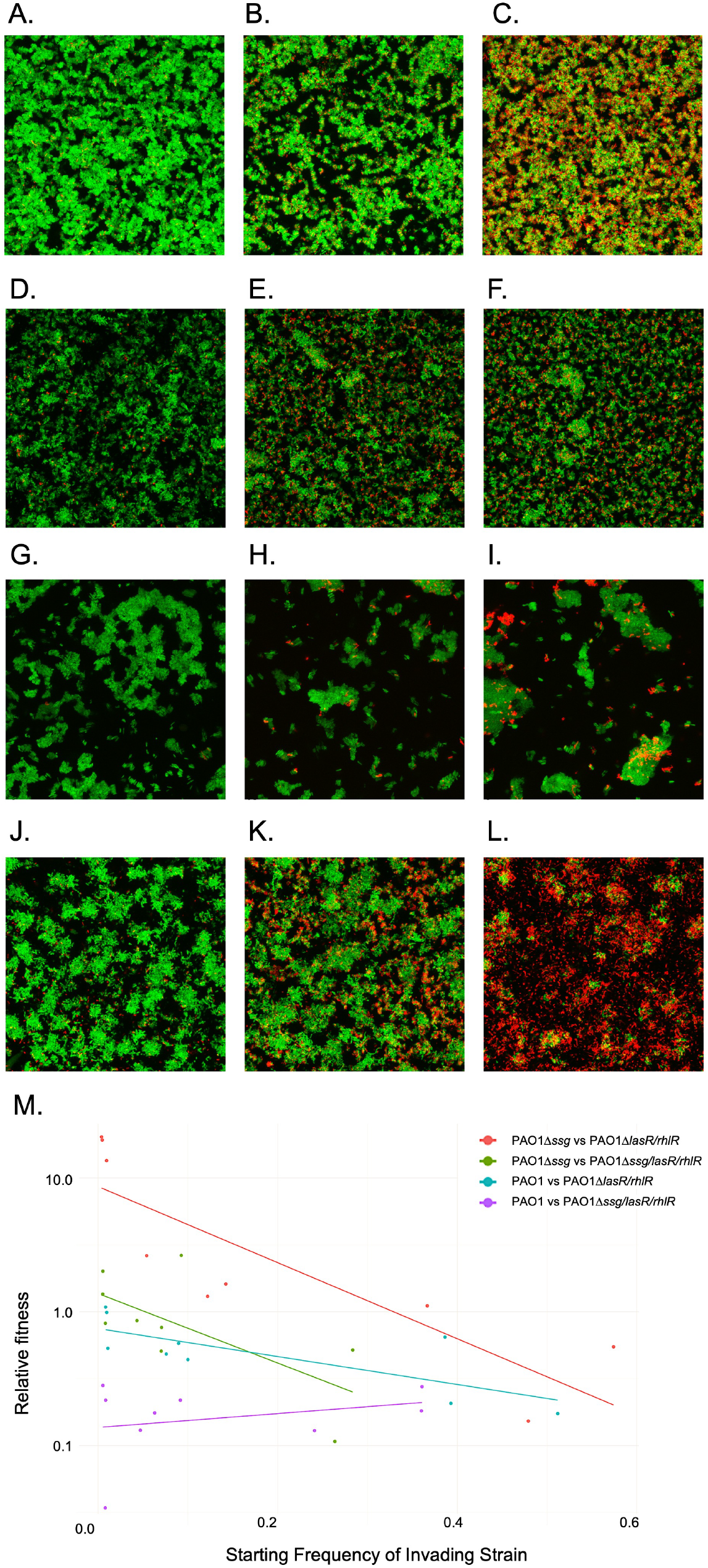
Proximity to cooperators enhances cheater relative fitness in social environments. Mixed populations of cooperators (PAO1, PAO1Δ*ssg*) and cheaters (PAO1Δ*lasR*Δ*rhlR*, PAO1Δ*ssg*Δ*lasR*Δ*rhlR*) were grown in QSM2 at three starting cooperator-to-cheater ratios of 99:1, 9:1, and 50:50. (A–L) Confocal images captured after 22 hours of growth for the following combinations: (A–C) PAO1::GFP + PAO1Δ*lasR*Δ*rhlR*::mCherry; (D–F) PAO1Δ*ssg*::GFP + PAO1Δ*ssg*Δ*lasR*Δ*rhlR*::mCherry; (G–I) PAO1::GFP + PAO1Δ*ssg*Δ*lasR*Δ*rhlR*::mCherry; and (J–L) PAO1Δ*ssg*:: GFP + PAO1Δ*lasR*Δ*rhlR* :: mCherry. (M) Cheater relative fitness at 24 hours plotted against starting frequency. Negative frequency-dependent fitness benefits were observed in all conditions except where hydrophobic cheaters were mixed with hydrophilic cooperators (G–I), the condition previously identified as having the highest cross-NNDist (least mixing).

In mixed populations of hydrophilic QS^+^ and QS^−^ cells, the distances between QS^−^ or QS^+^ cells were similar to distances between other cell types (mean NNDists for red, green and cross mean NNDists) and not significantly different (Fig 3I, Table S1-D). Similarly, in populations of hydrophobic QS^+^ and QS^−^ cells, there is no significant difference between distances between QS^−^ or QS^+^ and other cell types (mean NNDists for red, green and cross mean NNDists) (Fig. 3I, Table S1-D). In both mixed populations with homogenous cell surface properties, the highlighted area under the curve is small (Fig. S4B,C). The mean NNDist plots for hydrophobic-hydrophilic combinations of QS^+^ and QS^−^ cells reflected what was observed in the images, in a mixed population of hydrophilic QS^+^ and hydrophobic QS^-^ cells, the mean cross NNDist was significantly greater than red and green NNDists, with large fold changes (Fig. 3I, Table S1-D, p.adj < 0.05, foldchange = 4.20E-07 and 5.41E-12 respectively). In this mixed population, the red and green mean NNDists were the smallest yet observed in this analysis (-0.08 and -0.05 µm, respectively), while mean cross NNDist was the largest observed (0.51 µm). This pattern indicates hydrophilic QS^+^ cells spatially excluded hydrophobic QS^−^ cells, resulting in segregation between QS^+^ and QS-cells (Table S1-D). Additionally, the highlighted AUC was 82.64% (Fig. S4D). Similarly to asocial results, there was no highlighted AUC in hydrophobic QS^+^ cells mixed with hydrophilic QS^-^ cells (Fig. S4D), suggesting that hydrophilic QS^−^ cells can overcome differences in cell surface hydrophobicity and co-aggregate with hydrophobic QS^+^ cells.

### Liberation of Public Goods Through Proteinase K in QSM2 Restores Asocial Spatial Organization of Aggregates

We had some concerns that the social and asocial environments tested in these experiments have different salt bases, given that one is OS based, the other M9. We therefore sought a method to test the effects of public good availability without altering the salt base. While spiking proteinase K (PK) into wildtype cultures caused toxicity, it was possible to add 0.5 µg/mL PK into mixed QS^+^/QS^−^ populations in QSM2 to increase public good availability without causing cell death. Addition of PK confirmed that the change in spatial organization of aggregates seen in mixed populations of hydrophilic QS^+^ cells and hydrophobic QS^-^ cells was due to restrictions of social interactions and not due to the salt bases of media components (Fig. S4F-J, Table S1-E).

### Spatial Segregation Between QS^+^ and QS^−^ Cells Impacts the Fitness of QS^−^ Cells

As we determined that the mean NNDist between QS^+^ and QS^−^ cells was impacted by both the cell surface hydrophobicity and the social requirements of the environment, we next waned to determine whether there was a correlation between cell-to-cell proximity and the ability of QS^−^ cells to cheat on QS^+^ cells. We hypothesized that cross NNDist would correlate with cheater fitness. We seeded cooperators (QS^+^) and cheaters (QS^-^) at a ratio of 99:1, 9:1 or 50:50 and grew cells statically for 22-24 hours before imaging and cell counting (Fig. 4A-L). The non-cooperator cells were labeled with mCherry and a chromosomally inserted tn7-Trimethoprim marker. The mixed populations were plated on LBA for total cell counts and LBA-Tp500 for cheater counts at time 0 and after 24 hours. The relative fitness was calculated using the starting and ending ratios of cheaters, which was generated using the entire populations of three independent biological replicates (Fig. 4M).

The combination with the highest cross NNDist and lowest red and green NNDists (the treatment where cooperators and cheaters were farthest apart) was the only treatment where cheater fitness and starting concentration were not negatively correlated (hydrophilic cooperators and hydrophobic cheats; Fig. 4M, Table S2), normally referred to as “negative frequency-dependent fitness”. Additionally, out of all combinations, when a hydrophobic cheating strain was grown with a hydrophilic cooperator, it had the lowest fitness (as calculated by a linear model, 10:1 cooperator:cheat starting, Table S3). This indicated that the father apart cheats and cooperators are, the lower the relative fitness of the cheats. All other cheater-cooperator combinations resulted in negative frequency dependent fitness benefits for cheats (Fig. 4M, Table S2). Despite having similar density plots and NNDist means to the hydrophilic cooperator:cheat and hydrophobic cooperator:cheat mixed populations, hydrophilic cheats were significantly fitter than hydrophobic cheats in mixed cultures (Table S4). This indicates that the propensity for hydrophilic cells to cluster with themselves and exclude hydrophobic cells could be impacting directly on cheater fitness.

### Lower Fitness of Hydrophobic Cells Can Be Linked to Differences in Aggregate Spatial Patterns

Despite seeing separation between hydrophobic cooperators and hydrophilic cheats at the 1:10 mixed population, hydrophilic cheats had a high relative fitness (Fig. 4M), indicating that hydrophilic cheats are inherently fitter than hydrophobic cooperators in an environment where QS activation is required. We therefore tested invasion of a hydrophobic cooperator (PAO1Δ*ssg*) into a hydrophilic cooperator (PAO1) dominated population or vice-versa to determine whether both types of cooperator had a similar fitness when mixed. We mixed the cells 1:99, 1:9, 50:50, 9:1, 99:1 in QSM2 and plated at t=0, t=22 hours and imaged cells (Fig. S5A-F). We found that hydrophilic cooperators were inherently fitter than hydrophobic cooperators when mixed at all ratios (Fig. S5J). To confirm that hydrophobic cells are less fit than hydrophilic cells, we repeated our invasion experiment with a second hydrophobic variant: PAO1Δ*wbpL*. We found that similarly to competition with PAO1Δ*ssg*, PAO1 was able to increase in abundance against PAO1Δ*wbpL* at low starting frequencies in QSM2 (Fig. S5G-I).

## Discussion

Cooperative behaviors in bacteria, such as the shared production of public goods, are vulnerable to exploitation by non-producing cheats (5-8, 11, 24, 27, 28, 30, 53-60). Most explanations for how cooperative traits persist in microbes have focused on genetic or regulatory solutions, including kin recognition, metabolic constraints, or policing mechanisms (7, 28-30, 61-67). Our findings establish that the stability of social traits in *P. aeruginosa* is also fundamentally constrained by physiochemical cell surface properties governing aggregation and spatial organization. We previously showed that cell surface hydrophobicity plays a central role in organizing *P. aeruginosa* cells in polymer-rich and murine infection environments (42, 44). In this current study, we similarly found that hydrophilic wild-type cells formed loosely organized, stacked aggregates, whereas hydrophobic mutants lacking O-specific antigen (OSA) assembled into dense clumps. However, we also found that when mixed, hydrophilic and hydrophobic cells tended to segregate rather than intermix, creating an “oil-and-water” effect at the micron scale within aggregates.

This physical segregation had direct consequences for social interactions. QS cheaters were least successful when they were spatially separated from QS cooperators, indicating that access to shared resources was limited simply by distance and aggregate structure. Hydrophobic cheaters performed poorly when paired with hydrophilic cooperators, while hydrophilic cheaters gained a fitness advantage when interacting with hydrophobic cooperators. These outcomes suggest that the bacterial surface properties and aggregate architecture can effectively privatize public goods, shielding cooperators from exploitation without requiring active enforcement or additional metabolic costs. Our results therefore challenge the idea that the stability of cooperation is determined mainly by the energetic costs of cooperative traits (6, 8). Instead, they show that access shared resources often depend on how cells spatially arrange themselves. In this sense, biophysical constraints can set the stage on which, social evolution plays out, shaping the cellular interactions before genetic differences between strains come into effect.

Cell surface properties also influenced cell fitness beyond social cheating. Hydrophilic cells consistently outcompeted hydrophobic cells in mixed populations, even when QS activation and cheating was not required. This indicates that hydrophobic variants may carry an inherent disadvantage in aggregates. However, this barrier was not absolute. Under conditions where public goods were essential for growth, hydrophilic cheaters were able to grow close to and exploit hydrophobic cooperators, suggesting that strong metabolic growth incentives can partially overcome physical separation. In effect, dense clumped aggregates formed by hydrophobic cooperators may create steep local gradients of shared resources that diffuse to nearby cells, allowing limited exploitation at the aggregate boundary. Despite close spatial mixing between variants with similar surface properties, cheater fitness often remained relatively low. One possible explanation is the action of spiteful traits produced by cooperators, such as toxic secondary metabolites, which were not directly tested here (30, 68). Because cooperation, spite, and resistance to spite are often controlled by the same regulatory pathways such as quorum sensing (19), subtle variant differences could strongly influence outcomes. The short distances between cells with similar surface properties may make cheaters more vulnerable to such effects, whereas greater separation between dissimilar surfaces could reduce exposure. Future work examining how spatial structure interacts with spiteful behaviors would help clarify these dynamics.

Taken together, our findings underscore the central role of the physiochemical cell surface and aggregate structure in bacterial social evolution. Cell surface hydrophobicity shapes aggregate architecture, determines how closely cells associate, and ultimately governs access to shared resources and the success of cooperative or cheating strategies. These effects are likely to be especially relevant in vivo, particularly in the CF lung, where *P. aeruginosa* populations are genetically diverse, organized into spatially structured aggregates (40, 69-71), and frequently harbor mutations in QS and LPS biosynthesis pathways (45-48, 51, 72). Such heterogeneity is expected to promote the formation of distinct micron-scale niches that limit direct competition, stabilize coexistence, and modulate social interactions. More broadly, our results highlight the complex interplay between biophysical properties and social behaviors in spatially structured microbial communities. By demonstrating that cell surface properties and spatial proximity strongly influence aggregate organization and cheating success, this work extends current frameworks for understanding cooperation beyond genetic and metabolic costs alone. These insights have important implications for infection biology, as changes in spatial organization driven by surface and regulatory mutations are likely to alter microbial fitness, infection dynamics, and responses to treatment. Future studies should further explore how biophysical constraints interact with additional selective pressures, including cooperator-mediated spite, host immune responses, and antibiotic exposure, to shape the evolution and persistence of microbial populations in real-world environments.

## Materials and Methods

### Bacterial Strains and Growth Conditions

Strains used (with varying GFP, mCherry, Tn7-Tp or Tn7-Gm tags): PAO1, PAO1Δ*ssg*, PAO1Δ*lasR*Δ*rhlR*, PAO1Δ*ssg*Δ*lasR*Δ*rhlR* and PAO1Δ*wbpL*. To test social cheating in a spatially structured system, we developed three structured growth environments with differing social requirements. First, we used OS medium supplemented with 0.1% casamino acids (CAA) as a carbon source and 0.6 mg/mL eDNA to generate a structured media (OS-CAA) where *P. aeruginosa* forms aggregates. Mucin, used in SCFM2, was not required for aggregation in our model and was therefore excluded. This medium does not require QS activation for cellular growth and so we consider this an ‘asocial’ growth environment. To develop a “social” environment in which QS is essential for fitness, we developed a hybrid medium, by combining M9 salts with the metal components of OS medium: EDTA (8.56 mM), ZnSO_4_·7H_2_O (38.08 mM), FeSO_4_·7H_2_O (17.98 mM), MnCl_2_·4H_2_O (14.24 mM), CuSO_4_·5H_2_O (1.56 mM), CoCl_2_·6H_2_O (0.861 mM), H_3_BO_3_(1.89 mM), and NiCl_2_·6H_2_O (5.47 mM). We then added 1% BSA and 0.1% CAA to support QS activation and supplemented the medium with 0.6 mg/mL eDNA for spatial structure. This formulation was designated QSM2. Lastly, to generate an “asocial” version of QSM2 in which QS-regulated exoprotease activity was not required, we supplemented QSM2 with 0.5 μg/mL proteinase K (PK); higher concentrations led to toxicity for QS^+^ cells.

### Experimental Setup and Growth Conditions for Imaging

All experiments were performed in biological triplicate. Cheater strains were labeled with the plasmid pME6032:mCherry and a Tn7-Trimethoprim (Tn7-Tp) resistance marker, while cooperator strains were labeled with pME6032:GFP (73). Control experiments confirmed that swapping fluorescent tags did not influence the results. Plasmids were introduced by electroporation. Fluorescently labeled cells were grown overnight in 5 mL of Lysogeny Broth (LB) supplemented with 200 µg/mL tetracycline in 50 mL conical tubes at 37°C and 200 rpm. Cultures were washed twice with PBS, diluted 1:10 in PBS, and the optical density at 600 nm (OD_600_) was measured. We adjusted cell density to OD_600_ of 0.01 in 500 µL of the relevant media being used. When cooperator and cheater cells were mixed together, we used ratios of 99:1, 9:1, or 1:1. Media was prepared in bulk and aliquoted into microcentrifuge tubes. The appropriate cell volumes were added to reach 500 µL total per tube, mixed by vortexing, and 400 µL was transferred to a well of chamber slide (Lab-Tek) for imaging. The remaining 100 µL was serially diluted and used for colony-forming unit (CFU) counts to determine starting cell densities and verify inoculation ratios. Cultures were incubated statically at 37°C for 22 - 24 hours before confocal imaging and subsequent CFU enumeration.

### Cheater Fitness Quantification

To verify the initial frequencies of cheater cells, we plated a portion of the inoculum for colony-forming unit (CFU) enumeration. Twenty microliters of the starter cultures were serially diluted tenfold, four times, and plated on LB agar (LBA) for total CFU counts and on LBA supplemented with 500 µg/mL trimethoprim (LBA-Tp) for selective enumeration of cheaters. LBA plates were incubated at 37°C for 24 hours, and LBA-Tp plates were incubated for 48 hours to account for slower growth under antibiotic selection. After imaging, the entire 400 µL content of each Lab-Tek chamber well was transferred into a 2 mL screw-cap tube containing 5 mm metal beads. Wells were rinsed twice with 400 µL PBS, and the washes were added to each tube, yielding a total volume of 1.2 mL. Aggregates were disrupted using a bead beater, followed by vortexing at 14,800 rpm for 45 seconds. Samples were then serially diluted tenfold six times, and 50 µL aliquots were plated on LBA (total counts) and LBA-Tp (cheater counts). Plates were incubated at 37°C for 24 or 48 hours, respectively, before CFUs were enumerated. Experimental wells were plated for counts as described above. QS^−^ cell counts (growth on Tp500 plates) were divided by total cell counts (growth on LB plates) to determine starting and ending frequencies. Image analysis was not used to determine frequencies. Cheater fitness was calculated using LB plate counts for the whole population and LB Tp500 plate counts for cheater numbers and the following formula: v = (x_2_(1-x_1_))/(x_1_(1-x_2_)) where x_1_ is the initial proportion of mutants in the sample population and x_2_ is their final proportion (24). To determine if fitness was positively or negatively correlated with starting frequency, we performed a Pearson correlation using the cor function in RStudio. A linear model was fit to each treatment and used to predict the fitness of cheats at a starting frequency of 0.1. To predict the fitness of QS^−^ cells at a starting frequency of 0.1 pairwise comparison between different mixed populations was used by performing estimated marginal means using emmeans R package (74).

### Confocal Imaging and Image Analysis

After 22 hours of incubation, samples were imaged using a Zeiss Airyscan 880 confocal microscope set to Zoom 1 (scan area 134.95 µm × 134.95 µm). For each biological replicate, two 10 µm-thick Z-stacks were acquired using a 63× oil-immersion objective. Laser intensity and detector gain were optimized for each sample to achieve consistent image quality. All images were saved as .czi files for downstream analysis. Image analysis was performed using the MicrobeJ plugin in Fiji (ImageJ2). For both green and red fluorescence channels, we extracted x, y, and z coordinates of individual cells using the Batch processing feature with the following settings: “MultiThreading” under Experiment, “Channel 1” and “Dark” under Bacteria, and “Morphology 1 [Medial Axis]” under Morphology. The resulting coordinate data were exported as .csv files for spatial analysis. Spatial analysis was conducted in RStudio using the Spatstat package (75), which supports three-dimensional point pattern analysis. Separate coordinate files for the red and green channels were imported, and an empty matrix corresponding to the image dimensions (134.95 × 134.95 × 10 µm^3^) was generated. Nearest-neighbor distance measurements (NNDist) were calculated for same-color and opposite-color cell pairs using the nndist and nncross functions. The NNDist was calculated for each cell as the Euclidean distance to its closest neighbor within the specified environment. To quantify the spatial organization and spatial proximity of different cell types, the area under the curve (AUC) of measured NNDist distribution was calculated. The gray shaded region presented in plots indicating the portion of the cross-color (red–green) NNDist distribution where the AUC exceeds that of the same-color NNDist distributions. These gray regions are reported as a percentage of the total AUC of the red–green curve in each panel. Density plots of raw nearest-neighbor distance (NNDist) measurements for all cells pooled across images within each treatment. NNDist values were binned at 0.1 μm intervals. Curves represent distances between cells of the same color (green–green, green line; red–red, red line) and between differently labeled cells (red–green, black line). Raw NNDist data were Log10 transformed and outliers were removed, using a 1.5 interquartile range cut-off. Log10 transformed NNDist data were plotted as bar and whisker plots.

### Proteinase K Titration

To test the effects of the addition of an exogenous “public good”, we tested a titration of proteinase K titration. We grew PAO1::GFP, PAO1Δ*ssg*::GFP, PAO1Δ*lasR*Δ*rhlR*::mCherry Tn7-Tp, and PAO1Δ*ssg*Δ*lasR*Δ*rhlR*:: mCherry Tn7-Tp strains in biological triplicate in LB broth supplemented with 200 µg/mL tetracycline overnight. After washing as described above, we aliquoted cells into 96-well plates containing 100 µL of QSM2 and incubated them statically at 37°C for 24 hours. We measured OD600 using a plate reader (BioTek Synergy H1). We performed titrations of PK by preparing two-fold serial dilutions from 2 µg/mL to 0.0625 µg/mL. A t-test with Holm adjustment for multiple comparisons was run to compare QS^−^ growth to wildtype cells. All experiments performed in six technical replicates.

### Data visualization and statistical analysis

We used R v4.5.0 for all statistical analyses and data visualization using ggplot2 package. All codes are available on our GitHub (https://github.com/microlabazimi-collab/Biophysical-Constraints-Dictate-the-Stability-of-Social-Traits-in-Pseudomonas-aeruginosa-Aggregates).

## Acknowledgements

For funding, we thank the Georgia Institute of Technology and the National Institutes of Health for a grant (NIAID R56AI184449; R01AI153116) to S.*P*.D. We thank Cystic Fibrosis Foundation for a Fellowship (AZIMI18F0) to S.A and Georgia State University for startup funds to S.A. This work was additionally supported by the National Institute of Arthritis and Musculoskeletal and Skin Diseases of the National Institute of Health under (T32 AR056950-Musculoskeletal Research Training Program). Any opinion, findings, and conclusions or recommendations expressed in this material are those of the authors and do not necessarily reflect the views of the funding agencies.

## Author contributions

All authors contributed to research design; K.O.C. performed research and analyzed data; all authors contributed to writing the paper.

## Supplemental Material

**Supplemental Figure S1.**
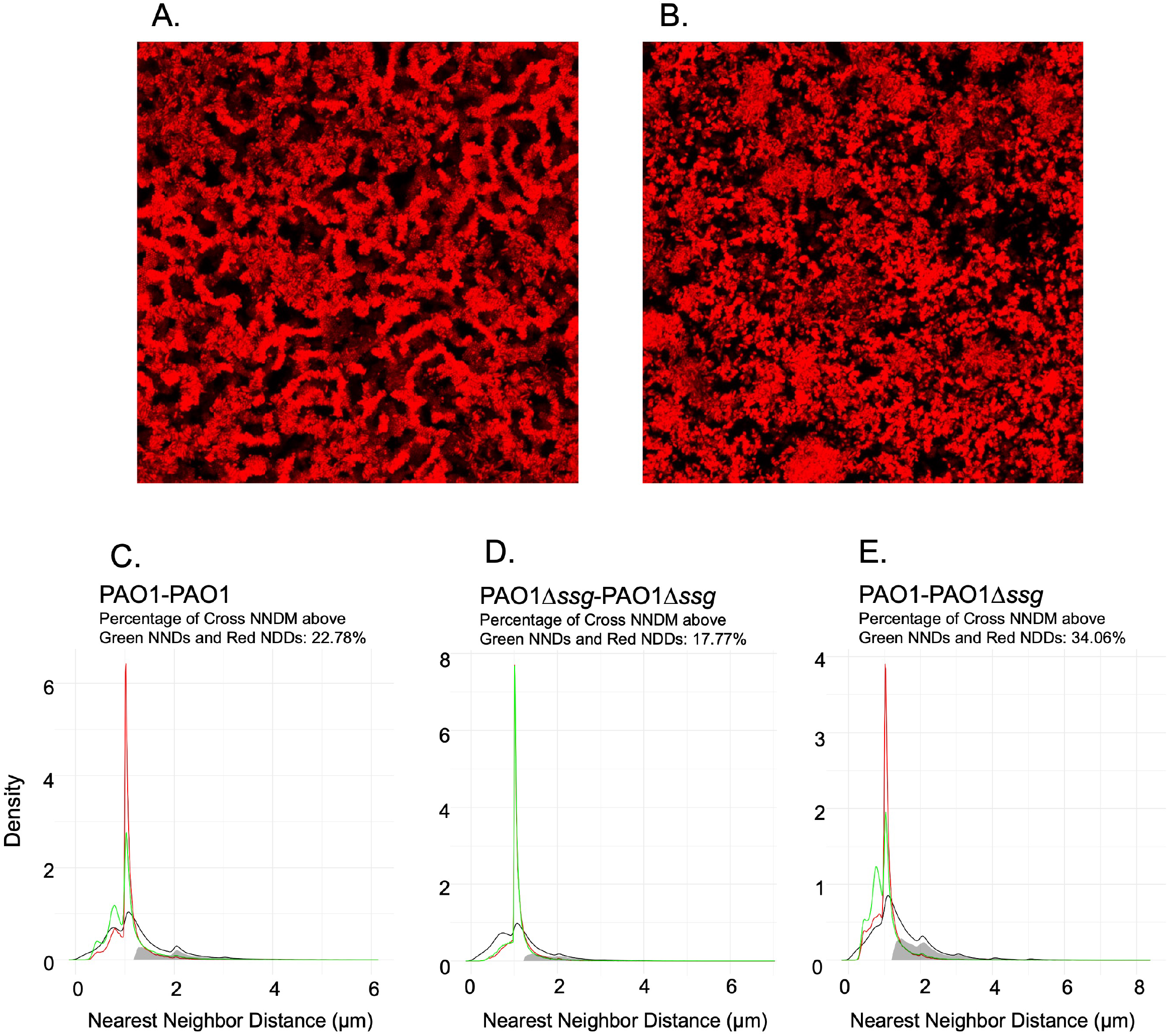
Label-flipping control in wildtype cells and raw nearest-neighbor distance (NNDist) density plots in OS-CAA. (A–B) Representative images showing that *P. aeruginosa* wildtype cells form stacked or clumped aggregates independent of fluorescent label identity (green or red). (C– E) Across all conditions, most NNDist measurements fall between ∼1–2 μm, indicating that neighboring cells are typically separated by this distance. The gray shaded region indicates the portion of the cross-color (red– green) distribution where the area under the curve (AUC) exceeds that of the same-color distributions. This gray region is reported as a percentage of the total AUC of the red–green curve in each panel title. Among the three tested conditions, the mixed population of hydrophobic and hydrophilic cells (E) shows the largest gray fraction (34.06%). This indicates a higher frequency of larger red–green NNDist values compared with red–red or green–green distances, suggesting that cells with different surface properties are more spatially separated from each other.

**Supplemental Figure S2.**
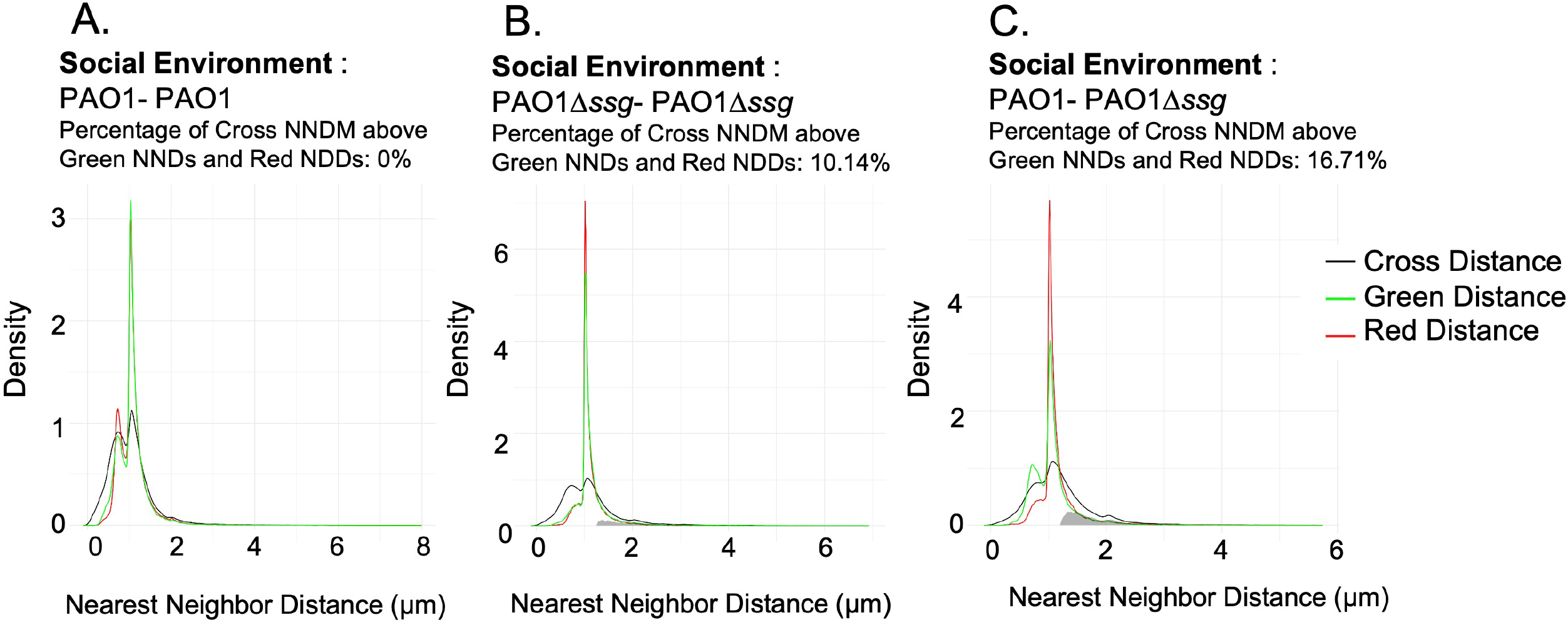
Raw nearest-neighbor distance (NNDist) density plots for wildtype cells in QSM2. (A–C) Density plots of raw NNDist measurements for red–red (red line), green–green (green line), and cross red–green (black line) cell pairs. NNDist values were pooled across all images for each treatment and plotted as density curves. The gray shaded region represents the portion of the cross-color (red– green) distribution where the AUC exceeds that of the same-color distributions (red–red or green–green). This gray area is expressed as a percentage of the total AUC of the red–green curve and is reported in the panel titles. Consistent with results observed in OS-CAA, the mixed population of hydrophobic and hydrophilic cells\ (C) shows the largest cross-distance enrichment (16.71%). This indicates a higher frequency of larger red–green NNDist values relative to same-color distances, suggesting increased spatial separation between cells with different surface properties.

**Supplemental Figure S3.**
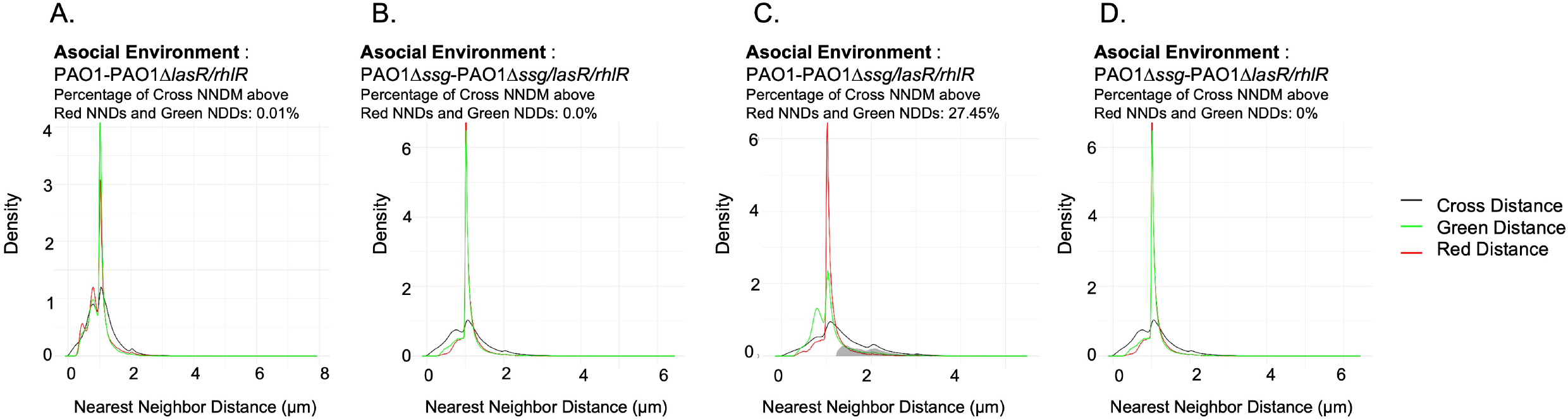
QS^+^ and QS^-^ cell combinations grown in OS-CAA and raw nearest-neighbor distance (NNDist) density plots. (A–D) Density plots of raw NNDist measurements for quorum-sensing proficient (QS^+^) and quorum-sensing deficient (QS^−^) cell combinations grown in OS-CAA. Curves represent distances between same-color cells (red–red, red line; green–green, green line) and between differently labeled cells (red–green, black line). NNDist values were pooled across all images within each treatment and plotted as density curves. The gray shaded region indicates the portion of the cross-color (red– green) distribution where the area under the curve (AUC) exceeds that of the same-color distributions (red–red or green–green). This gray area is expressed as a percentage of the total AUC of the red–green curve and is reported in the figure titles for each panel.

**Supplemental Figure S4.**
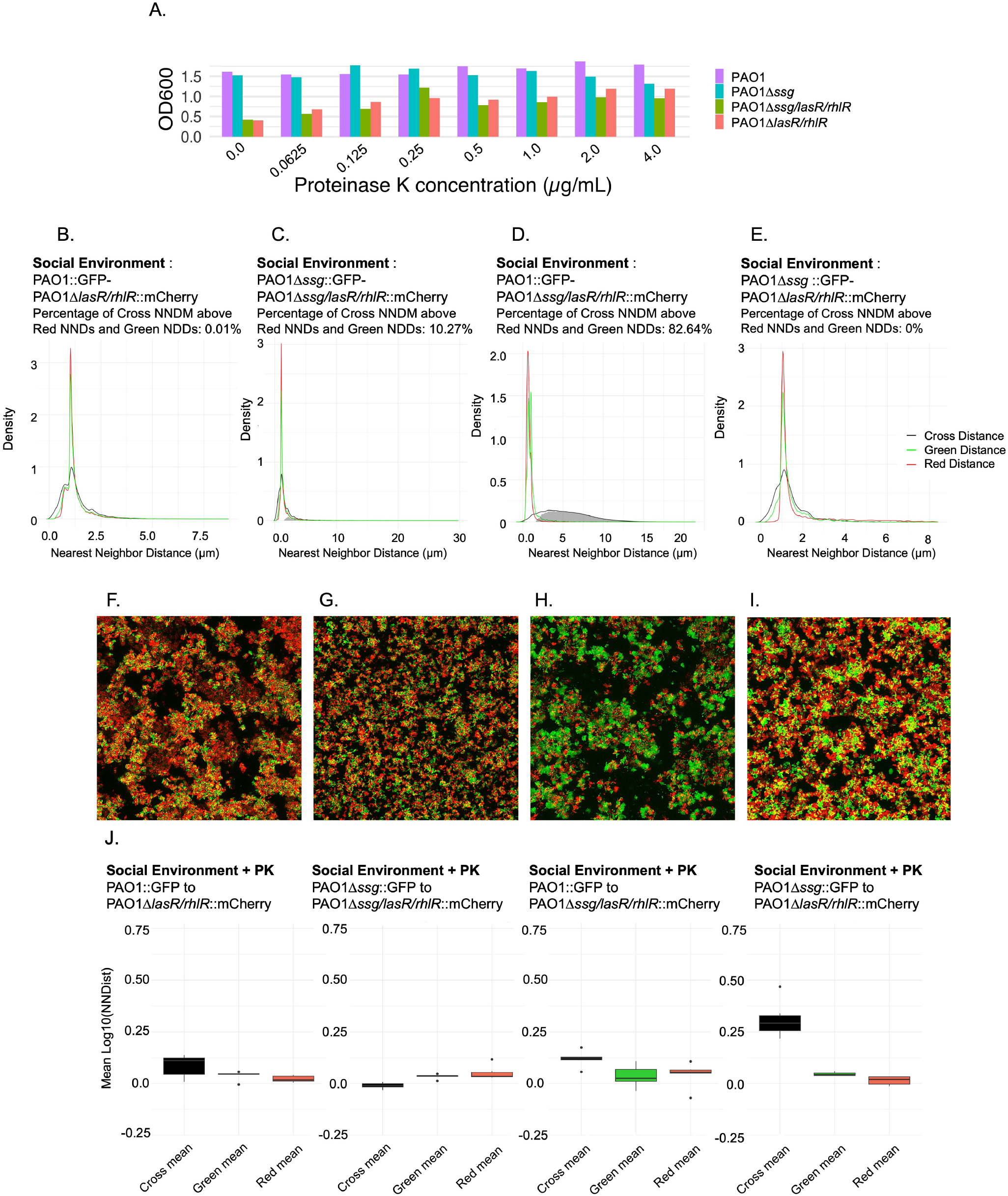
Effect of exogenous Proteinase K on QS^+^/QS^−^ interactions and nearest-neighbor distance (NNDist) distributions in QSM2. (A) Growth of quorum-sensing–deficient (QS^−^) strains in QSM2 supplemented with increasing concentrations of Proteinase K (PK). PK provides exogenous public goods by degrading protein substrates in the medium. Addition of PK increased the growth of both QS^-^ strains in a density-dependent manner. Statistical comparisons between QS mutant and wildtype growth were performed using t-tests with Holm correction for multiple comparisons. (B–E) Density plots of raw nearest-neighbor distance (NNDist) measurements for QS^+^/QS^−^ strain combinations grown in QSM2. Curves represent distances between same-color cells (red–red, red line; green–green, green line) and cross-labeled cells (red– green, black line). NNDist values were pooled across all images for each treatment and plotted as density curves. The gray shaded region indicates the portion of the cross-color distribution where the area under the curve (AUC) exceeds that of the same-color distributions; this value is reported as a percentage of the total AUC of the red–green curve in the panel titles. (F–I) Representative images of mixed QS^+^/QS^−^ populations grown in QSM2 supplemented with PK (0.5 µg/mL). (J–N) Boxplots showing red–red, green–green, and red– green NNDist measurements for QS^+^/QS^−^ mixtures grown in QSM2 + PK (0.5 µg/mL). For mixtures of cells with similar surface properties (hydrophobic–hydrophobic or hydrophilic–hydrophilic), cross NNDist values were generally not significantly different from same-color NNDists, with the exception of the PAO1::GFP + PAO1Δ*lasR*Δ*rhlR*::mCherry comparison. In contrast, mixtures of hydrophobic and hydrophilic cells showed significantly greater cross NNDists than red–red or green–green distances. The resulting NNDist distributions more closely resembled those observed in asocial (OS-CAA) medium than in social (QSM2) medium, indicating increased spatial separation between cells with different surface properties when exogenous public goods are supplied.

**Supplemental Figure S5.**
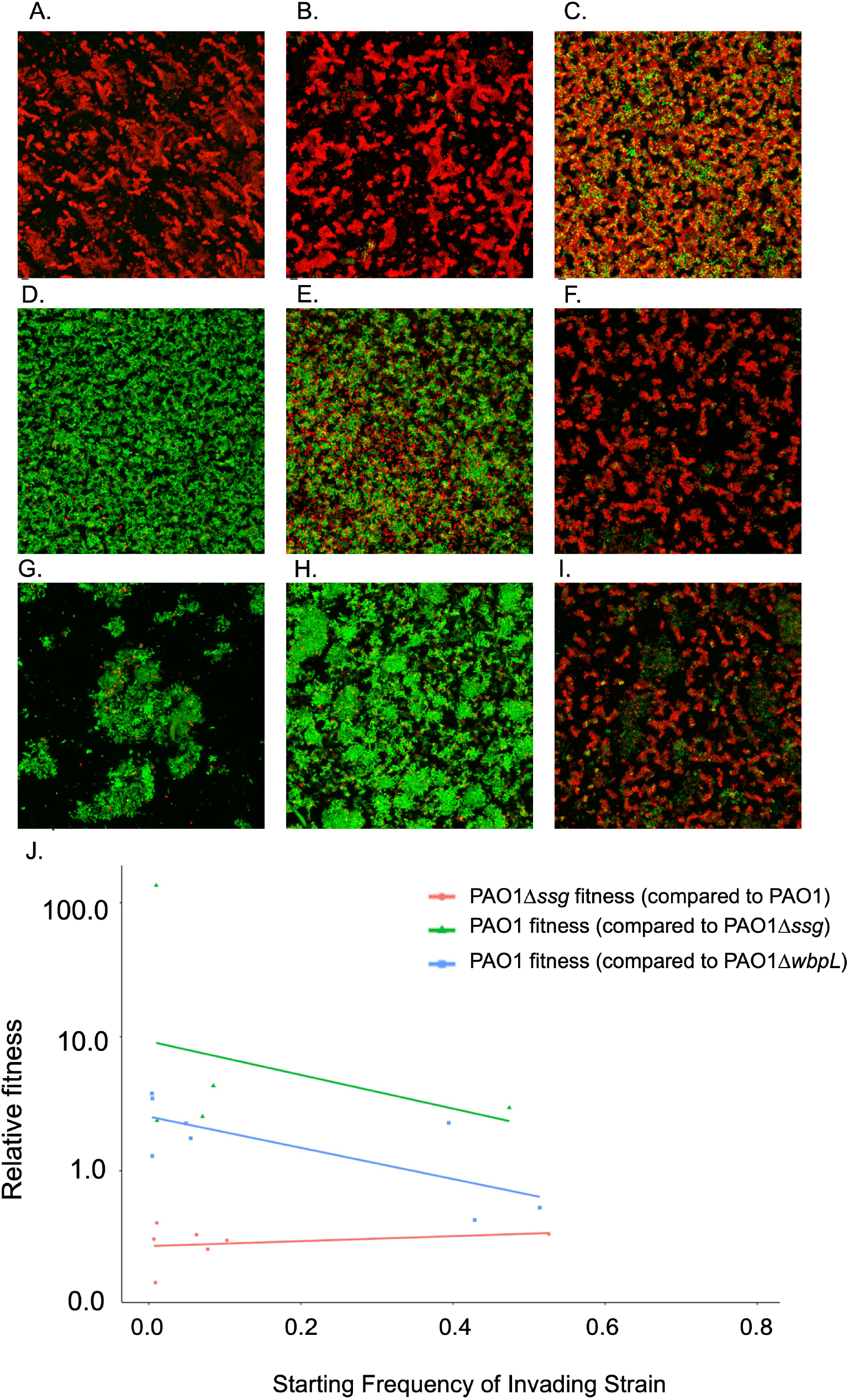
Hydrophilic cells outcompete hydrophobic cells in mixed populations. To determine whether hydrophilic cells have an inherent fitness advantage over hydrophobic cells, competition experiments were performed between PAO1::mCherry carrying a Tn7–trimethoprim resistance marker (Tp) and PAO1Δ*ssg* GFP carrying a Tn7–gentamicin resistance marker (Gm). These markers enabled tracking of strain frequencies during competition. Cells were mixed in QSM2 at starting ratios of PAO1:PAO1Δ*ssg* of 99:1 (A), 9:1 (B), 50:50 (C, F), 1:9 (D), and 1:99 (E), with a starting OD□□□ of 0.01. Cultures were allowed to grow approximately 100-fold before imaging and analysis. Initial cell densities and final densities after 22 hours of incubation were confirmed experimentally. Population sizes were quantified by enumerating the colony-forming units (CFUs) on LBA for total population size, LBA supplemented with 500 µg/mL trimethoprim (LBA-Tp500) to enumerate PAO1, and LBA supplemented with 200 µg/mL gentamicin (LBA-Gm200) to enumerate PAO1Δ*ssg*. PAO1 successfully invaded PAO1Δ*ssg* populations in a frequency-dependent manner and exhibited the highest relative fitness when initially rare. In contrast, PAO1Δ*ssg* was unable to invade PAO1 populations at any starting frequency. We additionally tested invasion of PAO1 mCherry Tn7-Tp into PAO1Δ*wbpL*::GFP populations, which lack OSA and form clumped aggregates. As observed with PAO1Δ*ssg* competitions, PAO1 displayed a frequency-dependent fitness advantage when competing against PAO1Δ*wbpL*. However, the magnitude of this advantage was smaller than that observed against PAO1Δ*ssg*.

## Table Legends

**Table S1: Normalized means of nearest neighbor distance measurement (NNDist) and comparisons**. Normalized NNDist in all tested mixed populations in asocial and social environments. Holms adjusted p-value.

**Table S2: Correlation between non-cooperator starting frequency and cheater non-cooperator fitness**. The relationship between non-cooperator starting frequency and non-cooperator fitness was examined to assess whether QS^−^ cells gain a negative frequency-dependent fitness benefit. In two out of four combinations there is significant negative correlation between QS^−^ cells frequency and their relative fitness (p < 0.05).

**Table S3: The fitness of cheats at a starting ratio of 10:1 as calculated by a linear model**. A linear model was built on the fitness data for each cheater-cooperator combination and used to determine the estimated fitness of cheats at a starting ratio of 10:1 cooperator:cheater. The higher the fit, the higher the estimated fitness of the cheater at that starting ratio.

**Table S4: Cheater fitness differences between cheater-cooperator combinations**. Linear models were built on the fitness data for each cheater-cooperator combination and used to determine the estimated fitness of cheats at a starting ratio of 10:1 cooperator: cheater; the cheater fitness difference between treatments at 9:1 starting frequency was calculated.

