## Supplemental tables for "Biophysical Constraints Dictate the Stability of Social Traits in *Pseudomonas aeruginosa* Aggregates"

**Supplementary Tables & Table Legends**

**Table Titles and Legends**

**Table S1: Normalized means of nearest neighbor distance measurement (NNDist) and comparisons.** Normalized NNDist in all tested mixed populations in asocial and social environments.

Holms adjusted p-value.

Table S1-A

| **Asocial** | Mean NNDist | | | Adj p-value | Fold Change | Adj p-value | Fold Change |
| --- | --- | --- | --- | --- | --- | --- | --- |
|  | Cross | Red-to-Red | Green-to-Green | Cross vs Red | Cross vs Red | Cross vs Green | Green vs Cross |
| **Mixed Populations** |  |  |  |  |  |  |  |
| PAO1 :: GFP - PAO1 :: mCherry | 0.08 | 0.02 | -0.01 | 0.10 | 4.01E+04 | 0.04 | 6.19E-12 |
| PAO1Δ*ssg* :: GFP - PAO1Δ*ssg* :: mCherry | 0.01 | 0.03 | 0.02 | 0.64 | 3.57E+00 | 0.61 | 4.11E+00 |
| PAO1 :: GFP - PAO1Δ*ssg* :: mCherry | 0.12 | 0.01 | -0.03 | 0.00 | 1.53E+09 | 0.00 | 2.85E-05 |

Table S1-B

| **Social** | Mean NNDist | | | Adj p-value | Fold Change | Adj p-value | Fold Change |
| --- | --- | --- | --- | --- | --- | --- | --- |
|  | Cross | Red-to-Red | Green-to-Green | Cross vs Red | Cross vs Red | Cross vs Green | Green vs Cross |
| **Mixed Populations** |  |  |  |  |  |  |  |
| PAO1 :: GFP - PAO1 :: mCherry | -0.05 | 0.01 | 0.00 | 1.00 | 1.36E-06 | 1.00 | 6.41E-44 |
| PAO1Δ*ssg* :: GFP- PAO1Δ*ssg* :: mCherry | -0.01 | 0.03 | 0.03 | 1.00 | 4.88E-01 | 1.00 | 5.27E-01 |
| PAO1 :: GFP- PAO1Δ*ssg* :: mCherry | 0.04 | 0.03 | 0.00 | 0.47 | 1.34E+01 | 0.00 | 6.91E-11 |

Table S1-C

| **Asocial Environment** | Mean NNDist | | | Adj p-value | Fold Change | Adj p-value | Fold Change |
| --- | --- | --- | --- | --- | --- | --- | --- |
|  | Cross | Red-to-Red | Green-to-Green | Cross vs Red | | Cross vs Green | |
| **Mixed Populations** |  |  |  |  |  |  |  |
| PAO1 :: GFP-PAO1Δ*lasR*/*rhlR* :: mCherry | 0.00 | -0.03 | -0.02 | 0.01 | 1.30E+00 | 0.03 | 1.48E+00 |
| PAO1Δ*ssg* :: GFP-PAO1Δ*ssg*/*lasR*/*rhlR* :: mCherry | 0.00 | 0.03 | 0.02 | 0.96 | 1.09E+00 | 0.94 | 1.10E+00 |
| PAO1 :: GFP-PAO1Δ*ssg*/*lasR*/*rhlR* :: mCherry | 0.09 | 0.03 | -0.01 | 0.00 | 1.48E+03 | 0.00 | 1.57E-09 |
| PAO1Δ*ssg* :: GFP-PAO1Δ*lasR*/*rhlR* :: mCherry | 0.19 | -0.02 | 0.00 | 0.00 | 1.65E-09 | 0.00 | 0.00E+00 |

Table S1-D

| **Social Environment** | Mean NNDist | | | Adj p-value | Fold Change | Adj p-value | Fold Change |
| --- | --- | --- | --- | --- | --- | --- | --- |
|  | Cross | Red-to-Red | Green-to-Green | Cross vs Red | | Cross vs Green | |
| **Mixed Populations** |  |  |  |  |  |  |  |
| PAO1 :: GFP-PAO1Δ*lasR*/*rhlR* :: mCherry | 0.08 | 0.08 | 0.08 | 1.00 | 9.49E+00 | 0.95 | 1.10E+01 |
| PAO1Δ*ssg* :: GFP-PAO1Δ*ssg*/*lasR*/*rhlR* :: mCherry | 0.14 | 0.10 | 0.08 | 0.90 | 2.64E+01 | 0.64 | 6.04E+01 |
| PAO1 :: GFP-  PAO1Δ*ssg*/*lasR*/*rhlR* :: mCherry | 0.51 | -0.08 | -0.05 | 0.00 | 4.20E-07 | 0.00 | 5.41E-12 |
| PAO1Δ*ssg* :: GFP-PAO1Δ*lasR*/*rhlR* :: mCherry | 0.06 | 0.09 | 0.07 | 1.00 | 5.17E+00 | 0.95 | 7.78E+00 |

Table S1-E

| **PK addition** | Mean NNDist | | | Adj p-value | Fold Change | Adj p-value | Fold Change |
| --- | --- | --- | --- | --- | --- | --- | --- |
|  | Cross | Red-to-Red | Green-to-Green | Cross vs Red | | Cross vs Green | |
| **Mixed Populations** |  |  |  |  |  |  |  |
| PAO1 :: GFP-PAO1Δ*lasRrhlR* :: mCherry | 0.09 | 0.02 | 0.04 | 0.04 | 1.57E+04 | 0.10 | 1.74E+02 |
| PAO1Δ*ssg* :: GFP-PAO1Δ*ssg*/*lasR*/*rhlR* :: mCherry | -0.01 | 0.05 | 0.03 | 1.00 | 6.05E-01 | 1.00 | 4.61E-01 |
| PAO1 :: GFP-PAO1Δ*ssg*/*lasR*/*rhlR* :: mCherry | 0.12 | 0.04 | 0.03 | 0.04 | 5.53E+02 | 0.02 | 2.87E+03 |
| PAO1Δ*ssg* :: GFP-PAO1Δ*lasR*/*rhlR* :: mCherry | 0.31 | 0.01 | 0.04 | 0.00 | 1.57E+23 | 0.00 | 7.93E+06 |

**Table S2: Correlation between non-cooperator starting frequency and cheater non-cooperator fitness.** The relationship between non-cooperator starting frequency and non-cooperator fitness was examined to assess whether QS¯ cells gain a negative frequency-dependent fitness benefit. In two out of four combinations there is significant negative correlation between QS¯ cells frequency and their relative fitness (p < 0.05).

| Mixed populations | Correlation | p-value |
| --- | --- | --- |
| PAO1 : PAO1∆*lasR/rhlR* | -0.675 | 0.0461 |
| PAO1 : PAO1∆*ssg*∆*lasR*∆*rhlR* | 0.219 | 0.571 |
| PAO1∆*ssg* : PAO1∆*lasR*∆*rhlR* | -0.682 | 0.0432 |
| PAO1∆*ssg* : PAO1∆*ssg*∆*lasR*∆*rhlR* | -0.474 | 0.197 |

**Table S3: The fitness of cheats at a starting ratio of 10:1 as calculated by a linear model.** A linear model was built on the fitness data for each cheater-cooperator combination and used to determine the estimated fitness of cheats at a starting ratio of 10:1 cooperator:cheater. The higher the fit, the higher the estimated fitness of the cheater at that starting ratio.

| Populations | Cheater Starting ratio | fitness | lower | upper |
| --- | --- | --- | --- | --- |
| PAO1 vs PAO1*∆lasR/rhlR* | 0.1 | 0.6513362 | -1.803981 | 3.106654 |
| PAO1 vs PAO1∆*ssg∆lasR/rhlR* | 0.1 | 0.1791583 | -2.149967 | 2.508284 |
| PAO1∆*ssg* vs PAO1*∆lasR/rhlR* | 0.1 | 9.1850678 | 6.686933 | 11.683203 |
| PAO1∆*ssg* vs PAO1∆*ssg/lasR/rhlR* | 0.1 | 1.0421401 | -1.23428 | 3.318561 |

**Table S4: Cheater fitness differences between cheater-cooperator combinations.** Linear models were built on the fitness data for each cheater-cooperator combination and used to determine the estimated fitness of cheats at a starting ratio of 10:1 cooperator: cheater; the cheater fitness difference between treatments at 9:1 starting frequency was calculated.

| **contrast** | estimate | SE | df | t.ratio | p-value |
| --- | --- | --- | --- | --- | --- |
| PAO1∆*ssg* : PAO1∆*lasR/rhlR* vs PAO1∆*ssg* : PAO1∆*ssg/lasR/rhlR* | 8.143 | 1.65 | 28 | 4.935 | 0.0002 |
| PAO1∆*ssg* : PAO1*∆lasR/rhlR* vs PAO1 : PAO1∆*lasR/rhlR* | 8.534 | 1.71 | 28 | 4.991 | 0.0002 |
| PAO1∆*ssg* : PAO1∆*lasR/rhlR* vs PAO1 : PAO1∆*ssg∆lasR/rhlR* | 9.006 | 1.67 | 28 | 5.401 | 0.0001 |
| PAO1∆*ssg* : PAO1∆*ssg*/*lasR/rhlR* vs PAO1 : PAO1*∆lasR/rhlR* | 0.391 | 1.63 | 28 | 0.239 | 0.9951 |
| PAO1∆*ssg* : PAO1∆*ssg/lasR/rhlR* vs PAO1 : PAO1∆*ssg/lasR/rhlR* | 0.863 | 1.59 | 28 | 0.543 | 0.9477 |
| PAO1 : PAO1∆*lasR/rhlR* vs PAO1 : PAO1∆*ssg/lasR/rhlR* | 0.472 | 1.65 | 28 | 0.286 | 0.9917 |
